# The Dynamic Nature of Genetic Risk for Schizophrenia Within Genes Regulated by *FOXP1* During Neurodevelopment

**DOI:** 10.1101/2025.05.12.653444

**Authors:** Deema Ali, Gary Donohoe, Derek W. Morris

## Abstract

*FOXP1* (Forkhead-box protein P1) is a crucial transcription factor in neural development and is associated with schizophrenia (SCZ). *FOXP1*-regulated genes may contribute to genetic risk of SCZ and this may vary across different stages of neurodevelopment. We analyzed transcriptomic data from mouse and human models of *FOXP1* loss-of-function across prenatal and postnatal developmental stages, including neural stem cells from embryonic mice (E14.5) and human brain organoids (equivalent to second trimester), and cortical tissues from different mouse postnatal stages P0, P7, and P47. P0 in mice corresponds to the third trimester in humans, while P7 and P47 represent early childhood and adolescence, respectively. The effect of *FOXP1* disruption on gene expression in cortical tissues/cells was assayed using RNA-seq, including time-course and pairwise gene expression analysis. Linkage disequilibrium score regression assessed if *FOXP1*-regulated genes were enriched for SCZ heritability. Gene-set enrichment analysis investigated if *FOXP1*-regulated genes were enriched for SCZ-associated genes reported as differentially expressed in single cortical cell studies. SynGO analysis mapped *FOXP1*-regulated genes to synaptic locations and functions. *FOXP1*-regulated genes are enriched for SCZ heritability, with significant results for E14.5, P7 and P47 but not P0. The P7 gene-set showed the strongest enrichment for SCZ-associated genes from single cortical cell studies. *FOXP1*-regulated genes at both P7 and P47 were involved in multiple synaptic functions and were mainly enriched within glutamatergic excitatory neurons, with P47 also showing enrichment within GABAergic inhibitory neurons across regions of the postnatal cortex. Prenatal *FOXP1*-regulated genes were enriched in progenitor cells and also mapped to the synapse. Genetic risk for SCZ within *FOXP1*-regulated genes follows a dynamic trajectory across developmental stages, showing stronger effects at timepoints that map to early childhood, followed by adolescence and second trimester.

**Author Summary:** Schizophrenia is a complex disorder caused by many genes. While existing treatments can help manage some positive symptoms, such as hallucinations and delusions, they are not effective in treating other life-limiting symptom areas. As a result, individuals with schizophrenia continue to face significant challenges, including disability, unemployment, homelessness, and social isolation. Genome-wide association studies of schizophrenia have been effective at identifying individual SNPs and genes that contribute to these phenotypes but have struggled to immediately uncover the bigger picture of the underlying biology of the disorder. Here we take an individual gene associated with schizophrenia risk, *FOXP1*, which is an important regulator of brain development. Using functional genomics data from models where *FOXP1* has been disrupted, we identified sets of genes regulated by *FOXP1* across different developmental stages, focusing on the cortical region of the brain. Our findings reveal that *FOXP1*-regulated genes are most strongly associated with schizophrenia during early childhood and adolescence. These genes are associated with synaptic components, including presynaptic and postsynaptic structures. By integrating developmental timing, cell type specificity, and functional pathways, this study provides valuable insights into the molecular mechanisms underlying schizophrenia.

## Introduction

Forkhead Box P1 (*FOXP1*) belongs to the FOX family of transcription factors that coordinate essential developmental processes, including within the nervous system [1–3]. *FOXP1* is associated with a rare neurodevelopmental disorder (FOXP1 syndrome) [4–7]. where different types of mutations have been identified as causal [5,8–10]. Additionally, common variants in *FOXP1* have been associated with schizophrenia (SCZ) [11,12], general cognitive ability [13] and autistic spectrum disorder (ASD) [14].

*FOXP1* is expressed in both the developing and adult brain [3,15–17] and is a key regulatory gene in neural development, contributing to transcriptional mechanisms involved in neurogenesis, neuronal migration, morphogenesis, and synaptic plasticity [18–21]. Several studies have generated *FOXP1* knockout (KO) and knockdown (KD) models to investigate the functional role of *FOXP1* in the brain and to explore the molecular pathways underlying human phenotypes associated with *FOXP1*. In the early stages of development, high FOXP1 levels in apical radical glial cells (aRGCs) are associated with early neurogenesis in human cortical development, while lower levels are linked to later stages [21]. FOXP1 is also involved in basal radial glial cell (bRGC) formation, where its dysregulation impairs their proliferation and differentiation, causing a long-term reduction in the number of excitatory cortical neurons [21,22]. In later stages of development, *FOXP1* expression is predominantly confined to the pyramidal neurons of the neocortex. Deletion of *foxp1* using a conditional knockout (cKO) approach (Emx1.Cre; Foxp-1 flox/flox) has been reported to cause abnormalities in vocal communication as well as neocortical cytoarchitectonic alterations via neuronal positioning and migration at early postnatal stages [19]. Apart from cortical tissue, *FOXP1* is also expressed in the medium spiny neurons of the striatum as well as in the CA1/CA2 hippocampal subfields [3,23,24]. It plays a role in the differentiation of dopamine neurons in the midbrain and medium spiny neurons in the striatum [25,26]. Adult mice with brain specific homozygous deletion of *foxp1* demonstrated developmental abnormalities in the striatum and hippocampus, dispersed neuronal organization in hippocampal CA1, reduced excitability, larger excitatory postsynaptic current amplitudes in CA1 neurons, impaired short-term memory, and ASD-like behaviors [16].

Expression analysis in a rat SCZ model identified *FOXP1* as a novel SCZ candidate gene [27]. In a recent single-nuclei RNA sequencing (snRNA-seq) study of postmortem prefrontal cortical tissue, *FOXP1* was prioritized as one of the key TFs targeting SCZ-associated differentially expressed genes (DEGs) in neuronal populations [28]. Additionally, Levchenko *et al*., (2021) suggested that *FOXP1* may contribute to immune system alterations in SCZ through interactions with immune-related genes involved in NFκB-mediated inflammatory responses, which are upregulated in SCZ [29].

Here, we explore *FOXP1*’s contribution to SCZ using RNA-seq data from *FOXP1* loss-of-function models, focusing on its involvement in cortical development. Given that *FOXP1* is expressed in both developing and mature brains, and that SCZ is a disorder that is linked to brain development from in utero through to adulthood [30,31], we utilized data generated from prenatal and postanal stages, including cortical neural stem cells (NSCs) and bRGCs from timepoints that reflect the second trimester in humans, as well as data from mouse neocortical tissues at various developmental stages - P0 (birth; equivalent to third trimester in humans), P7 (7 days after birth; equivalent to early childhood in humans), and P47 (47 days after birth; equivalent to adolescence in humans) - to investigate the involvement of *FOXP1*-regulated genes in SCZ risk. These timepoints represent critical stages for the pathophysiology of neurodevelopmental disorders such as SCZ. We investigated whether *FOXP1-*regulated genes at various developmental stages are enriched for genes containing common SNPs associated with SCZ from GWAS and for genes differentially expressed in single cell types from the prefrontal cortical brain region of SCZ patients and control samples. Additionally, we investigated the impact of *FOXP1* disruption on synaptic processes and specific cell types to better understand how *FOXP1*-regulated genes contribute to the molecular mechanisms underlying SCZ. Finally, we aimed to identify trans-expression quantitative trait loci (trans-eQTL) at *FOXP1* that are associated with altered expression of *FOXP1* downstream target genes.

## Materials and Methods

### Ethics statement

Data were directly downloaded from published studies and no further ethics approval was required. Each study is referenced, and information about ethics approval is provided in the original references.

### Sourcing Transcriptomic Data for *FOXP1* Loss-of-Function Models

We analyzed transcriptomic data derived from studies that generated *FOXP1* loss-of-function models in both mouse and human across different developmental stages, including prenatal and postanal periods. For the prenatal stages, three data resources corresponding to the second trimester of human fetal development were used. The first included NSCs at embryonic day 14.5 (E14.5) of mouse development. In this study, RNA-seq was performed on isolated embryonic NSCs that were transduced *in vitro* with either a short hairpin RNA (shRNA) targeting *foxp1* for knockdown or a scrambled shRNA as a control (CTL), with two samples analyzed for each condition [18]. The second data source included bRGCs derived from human cerebral organoids [22]. CRISPR-Cas9 technology was used to induce *FOXP1*-KO in human induced pluripotent stem cells (hiPSCs), which were subsequently differentiated into cerebral organoids. snRNA-seq was then conducted on the cortical regions of the organoids to compare gene expression profiles of KO and CTL phenotypes, with three samples analyzed for each condition [22].The third source included mouse neocortical tissues at P0 (birth), which, although postnatal in mice, corresponds to the third trimester in humans and is therefore considered a prenatal stage for this study [32]. For postnatal stages, mouse neocortical tissues at P7, and P47 were analyzed [32,19], corresponding to early childhood, and adolescence stages of brain development in humans respectively. *foxp1*-KO mice were generated using a brain-specific conditional KO of *foxp1* via the Cre-Lox system. RNA-seq was subsequently performed on extracted RNA from the neocortical tissues of both *foxp1*-KO and wild-type (WT) controls mice at P0, P7, and P47, with four samples analyzed for each condition and developmental stage [19,32].

### Transcriptomic Analysis of *FOXP1* at Various Developmental Stage

RNA-seq raw data were downloaded directly from the Gene Expression Omnibus (GEO) database, with the following accession numbers corresponding to each dataset: GSE101633 for the embryonic NSCs, GSE98913 for neocortical tissues at P0 and P7, and GSE97181 for neocortical tissues at P47. FastQC (v0.12.1) (http://bioinformatics.babraham.ac.uk/projects/fastqc/) was used for quality assessment of reads. Raw reads were trimmed for adapters using Trimmomatic (v0.39) [33]. Filtered reads were then aligned to the mouse genome mm10 (https://genome.ucsc.edu) using HISAT2 (v2.2.1) [34]. The BAM alignment files were then subjected to featureCounts (v2.0.3) for read counting [35]. The list of DEGs from bRGCs was directly extracted from the original paper without further analysis [22].

### Time-Course Gene Expression Analysis

Time-course gene expression analysis was conducted to examine the differential effects of *foxp1*-KO across various developmental stages by applying a likelihood ratio test (LRT) using the DESeq2 R package [36,37]. This test compared the likelihood of the data under a full model (Stage + Condition + Stage:Condition) with that under a reduced model (Stage + Condition). Only data from the P0, P7, and P47 timepoints were included in the analysis. The earliest developmental stage, P0, was set as a reference stage and WT as a reference condition. The significant genes were identified at FDR < 0.05. Clustering analysis was conducted on the genes identified in LRT analysis to identify gene groups exhibiting specific expression patterns across KO and WT samples separately. The gene count data generated from pair-wise gene expression analysis (described below) was first subjected to regularized log (rlog) transformation. Clustering was then performed using the divisive hierarchical clustering method, implemented in the degPatterns function from the DEGreport R package [38]. Only genes identified from the time-course gene expression analysis with FDR of less than 0.05 were included in the clustering process.

### Pair-Wise Gene Expression Analysis

Pair-wise differential gene expressions between *foxp1*-KO and WT controls for each development stage were performed separately for each timepoint. Counts per million (CPM) values were calculated, and genes with values of 1.0 or higher in at least two replicates for either the KO or WT conditions were considered. DESeq2 (version 1.44.0) was used to detect the DEGs [36,37]. The significant DEGs were identified at false discovery rate (FDR) < 0.05. Gene annotation, the conversion of mouse genes to their human orthologs, was conducted on the transcripts using the BiomaRt package (version 2.60.1) [39,40].

### Stratified Linkage Disequilibrium Score Regression Analysis

Stratified linkage disequilibrium score regression (sLDSC) (https://github.com/bulik/ldsc) [41] was used to investigate if the *FOXP1* gene-sets across the different developmental stages were enriched for heritability contributing to SCZ. GWAS summary statistics for schizophrenia (SCZ; 76,755 cases and 243,649 controls) [12] was obtained from publicly available databases (the Psychiatric Genomics Consortium website; www.med.unc.edu/pgc). For control purposes, we carried out sLDSC analysis using GWAS summary statistics for an additional four brain-related disorders, including major depressive disorder (MDD) [42], obsessive–compulsive disorder (OCD) [43], Alzheimer’s disease (AD) [44] and stroke [45]. Linkage disequilibrium (LD) scores between SNPs were estimated using the 1000 Genomes Phase 3 European reference panel. SNPs present in HapMap 3 with an allele frequency > 0.05 were included. Heritability was stratified in a joint analysis between 53 previous function genomic annotations and each *FOXP1* gene-set. Enrichment for heritability was compared to the baseline model using the Z-score to derive a (one-tailed) P-value. A Bonferroni correction was applied to determine significant enrichments, which set the corrected P value threshold at < 1.11E-03.

### Competitive Gene-Set Analysis of *FOXP1* in SCZ

Competitive gene-set enrichment analysis (GSEA) using the R package, *fgsea* (Fast Gene Set Enrichment Analysis)[46] was used to test if the *FOXP1*gene-sets were enriched for cell-type specific DEGs for 29 different cell-types (S1 Table) derived from single-nuclei RNA sequencing (snRNA-seq) of the prefrontal cortical brain region comparing SCZ and control samples [28]. *fgsea* was conducted pre-ranked mode, where input gene-sets from DeSeq2 were ranked by the Wald statistic. A competitive comparison was then performed to determine whether genes that feature in a set are highly ranked in terms of differential expression compared to genes that are not in the set. Gene-sets with an FDR corrected p-value < 0.05 were considered significantly enriched.

### Cell-Type Enrichment Analysis of *FOXP1* Gene Sets

The Expression Weighted Cell-type Enrichment (EWCE) R package (https://github.com/NathanSkene/EWCE) was used to assess if the *FOXP1* gene-sets had higher expression in a particular cell type than expected by chance [47]. This method generates random gene sets (*n* = 100,000) of equal length from background genes to estimate the probability distribution. We performed enrichment analysis in a prenatal human dataset and in an adolescent mouse dataset [48,49]. The prenatal human dataset includes snRNA-seq data from three second-trimester fetuses and encompasses different brain regions. However, the analysis was restricted to 17 distinct clusters of nuclei from the frontal cortex (FC) [48]. The adolescent mouse dataset includes data from 19 regions across the central and peripheral nervous system of mice at post-natal days 12-30 and at 6- to 8-weeks [49]. The significance of the enriched expression of the *FOXP1* gene-sets relative to the background genes in each cell type was assessed by calculating the difference in standard deviations between the two expression profiles. The significant cell types were identified at FDR < 0.05.

### Functional Enrichment Analysis

To analyze the enrichment of synaptic gene ontologies among the *FOXP1* gene-sets, we used SynGO (https://www.syngoportal.org/) [50], an expert-curated resource for synaptic GO analysis.

Analyses for GO terms, including biological processes and cellular components were performed. The analysis used cortex tissue-expressed genes (n=16,985) as a background gene-set (S2 Table). Ontologies with an FDR corrected p-value < 0.01 were considered significantly enriched.

## Results

### Analysis of FOXP1 in Neocortical Tissues

#### Time-course Gene Expression Analysis

We first conducted time-course gene expression analysis to determine if the effect of *foxp1*-KO on downstream gene expression differed between any of the timepoints. This analysis identified 1,146 genes (FDR < 0.05; S3 Table) where the effect of *foxp1*-KO on their expression differed over time. Of these mouse genes, 1,065 have human orthologs. sLDSC analysis showed that these genes were significantly enriched for genes associated with SCZ (*P*=3.45E-04; S4 Table). To further explore these significant genes, their regularized log2-transformed counts were grouped based on similar expression patterns, resulting in the formation of 20 distinct clusters (S5 Table). The expression patterns across the developmental stages for each cluster, along with the scaled expression levels of individual genes, are illustrated in Fig 1. Among the identified genes, 35 are located with genome-wide significant loci for SCZ, and 7 of these are among the 120 genes prioritized using fine-mapping and functional genomic data (S6 Table). Additionally, 2 of the identified genes, *XPO7* and *DNM3*, are located within coding variants with FDR < 5% in the SCHEMA study [51]. A number of clusters (clusters 12-15 and 17-20) exhibited distinct expression patterns between the WT and KO groups across one or more developmental stages (Fig 1). Among the genes in these clusters, three (*PLK2*, *CACNA1I*, and *NEK1*) are located with genome-wide significant loci for SCZ. Overall, this analysis indicates that there is a dynamic effect where *foxp1*-KO can have different levels of impact on the expression of genes under its influence at different stages of development. It is therefore warranted to investigate the genes expressed at each timepoint for their contribution to SCZ. Fig 2 provides an illustration of the stepwise methods used from here to study *FOXP1*-regulated genes over time.

**Fig 1.**
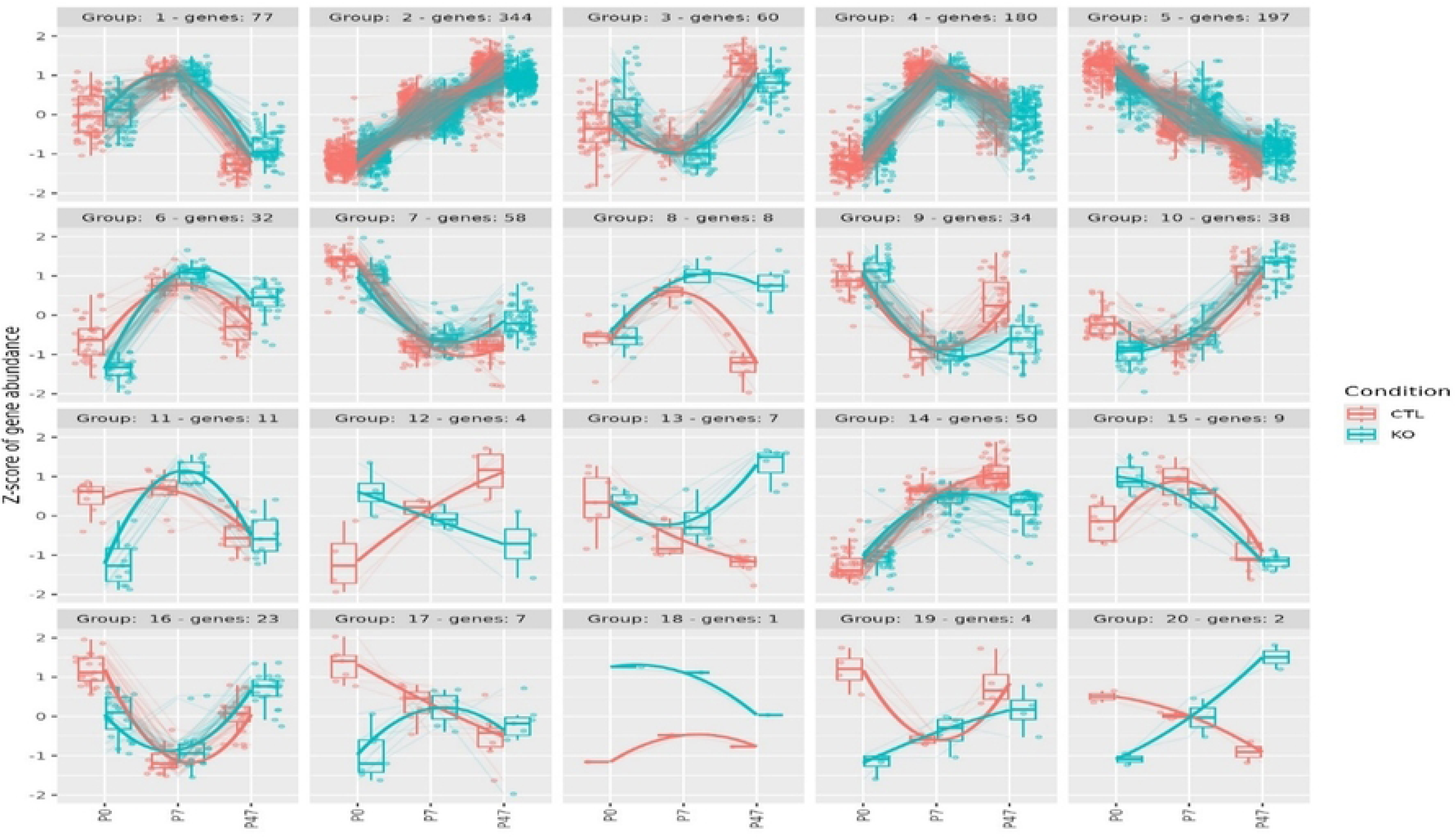
Clusters of Significant Genes Identified by Time-Course Gene Expression Analysis Across Different Developmental Stages in Mouse Neocortical Tissue. Divisive hierarchical clustering was performed for 1,146 genes (FDR < 0.05) according to log2 normalized read counts. The cluster number and the corresponding number of genes are provided for each cluster. Genes are plotted on the y-axis according to the scaled expression value (zscore). Lines visualize the expression pattern across development, connecting the average expression level at each stage for genes within each cluster. KO: Knockout; CTL: Control.

**Fig 2.**
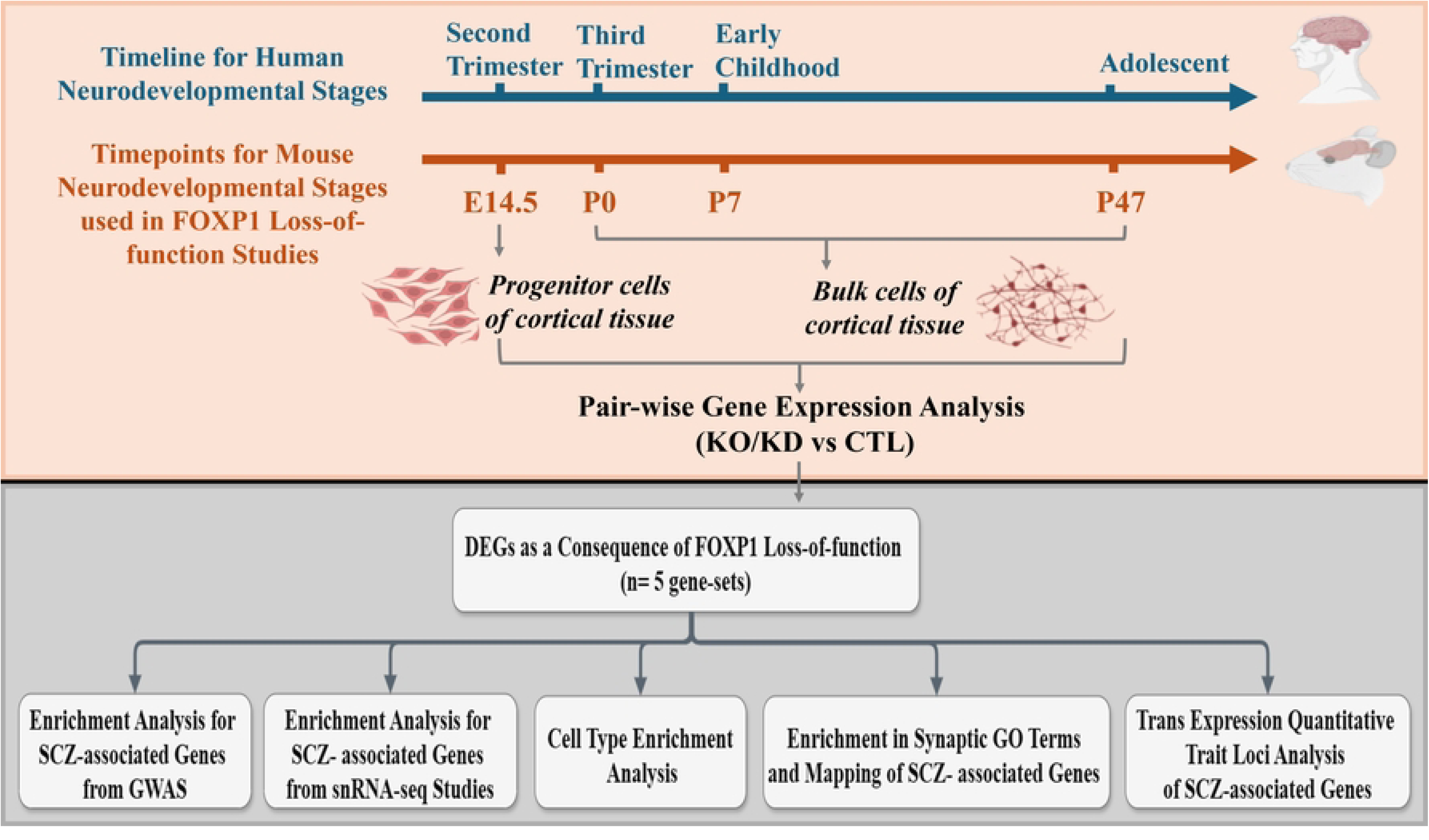
Schematic of Pairwise Gene Expression Analysis and Functional Enrichment Methodology for FOXP1 Loss-of-Function Models. E: embryonic; P: Postnatal day; KO: Knockout; KD: Knockdown; CTL: Control; DEGs: Differentially expressed genes; SCZ: Schizophrenia; GWAS: Genome-wide association study; snRNAseq: Single nucleus RNA sequencing.

### Pair-wise Gene Expression Analysis

We identified 423 DEGs (186 up-regulated and 237 down-regulated) at P0, 394 DEGs (139 up-regulated and 255 down-regulated) at P7 and 1,527 DEGs (712 up-regulated and 815 down-regulated) at P47 (Fig 3 and S7 Table). In total, there are 32 DEGs common to all three timepoints, with the vast majority showing concordant effects (i.e., 14 genes are down-regulated at each timepoint, and 15 genes are up-regulated at each timepoint). These data show that there is some overlap across stages but that the majority of DEGs are stage-specific, suggesting that *FOXP1* influences the expression of different genes at different stages during development.

**Fig 3.**
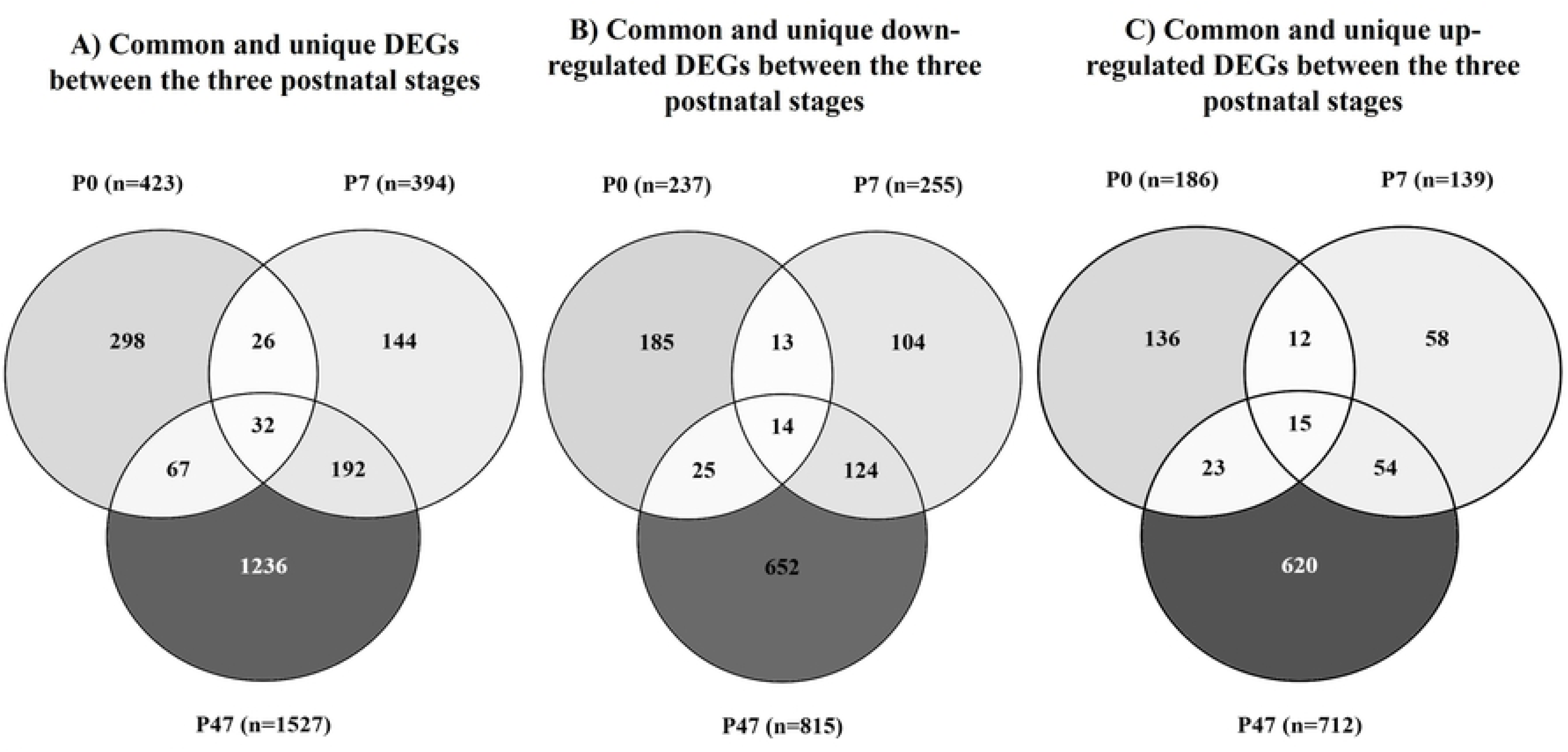
Overlap of DEGs between FOXP1 KO-vs-WT at the postnatal developmental stages. Venn diagrams of A) all DEGs, B) down-regulated DEGs, and C) upregulated DEGs. These Venn diagrams illustrate the number of shared and unique DEGs across postnatal developmental stages. P: Postnatal day; DEGs: Differentially expressed genes.

### Enrichment Analysis for Schizophrenia-associated Genes from GWAS

Using sLDSC regression analysis, we observed that there was no enrichment for genetic risk for SCZ among DEGs at P0 (*p*=0.105) but this quickly changed for P7 (*p*=4.9E-06) and P47 (*p*= 5.0E-08; Fig 4A andS4 Table). There is overlap between P7 and P47 (n=224) but even when excluding these from either gene-set, both remain significantly enriched (S4 Table). No significant enrichment was detected for any of the four control phenotypes (S4 Table). These findings highlight that the contribution of genes influenced by *FOXP1* to SCZ is developmental stage-specific; genes regulated by *FOXP1* at P0 contribute little to genetic risk for SCZ while just a short time later (P7), variation in *FOXP1*-regulated genes does contribute to genetic risk for SCZ and this is maintained at the later P47 timepoint, even though the set of DEGs changes considerably.

**Fig 4.**
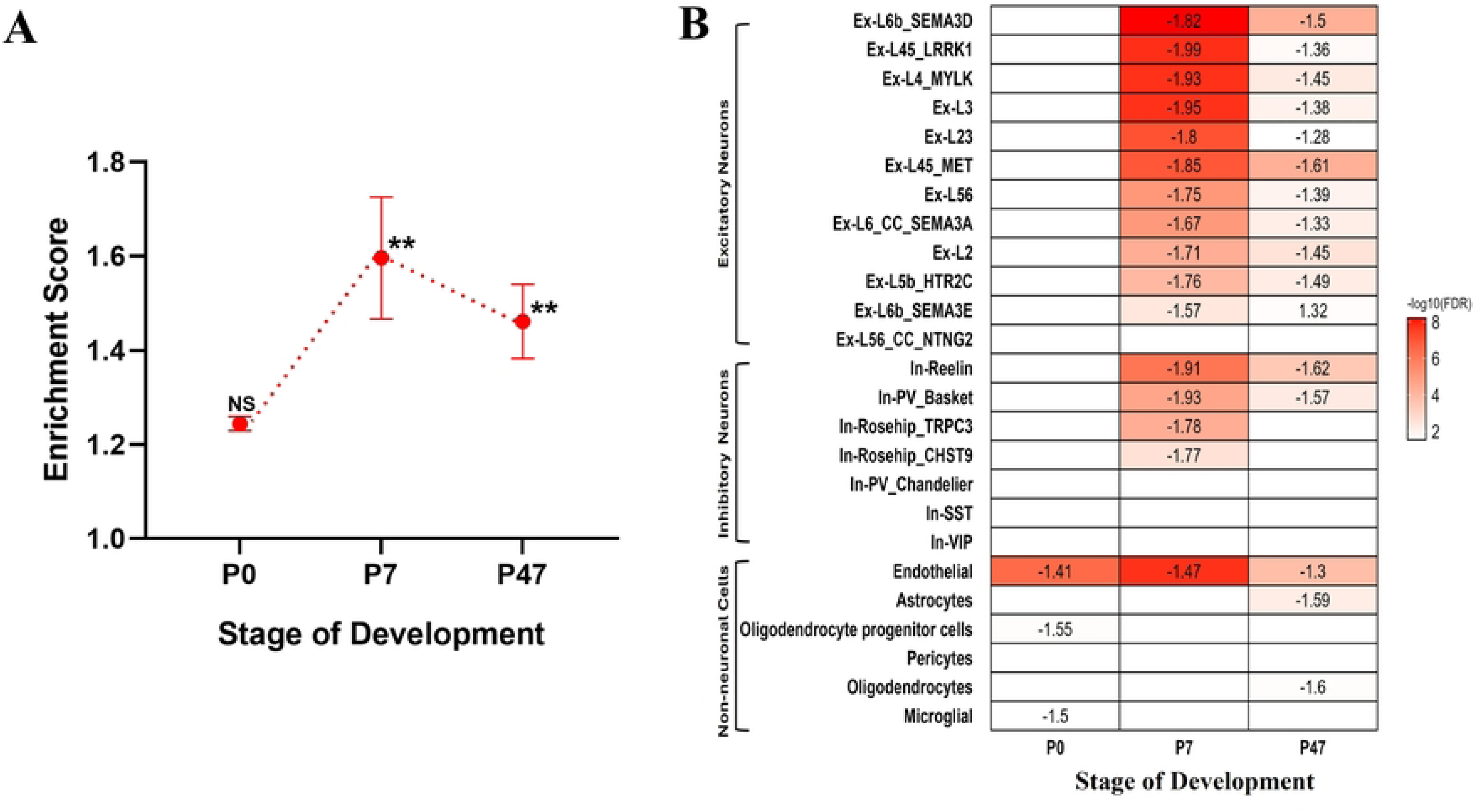
Enrichment of FOXP1 Gene-Sets in SCZ-Associated Genes Across Different Developmental Stages in mouse neocortical tissue. (A) Results from sLDSC analysis of FOXP1 gene-sets across the different developmental stages using SCZ-GWAS data. The graph plots the enrichment values, defined as the ratio of heritability (h2) to the number SNPs, on the y-axis. The x-axis represents the different developmental stages. Two asterisks (**) indicate significance after Bonferroni correction, one asterisk (*) indicates nominal significance, and “NS” indicates not significant (p>0.05). (B) Heatmap shows the enrichment of FOXP1 gene-sets in SCZ associated gene-sets across the different developmental stages based on snRNA-seq data. The intensity of color represents the –log (FDR), with darker colors indicating more significant enrichment. Normalized Enrichment Score (NES) values are shown in the cells. A significant negative NES value indicates that members of the gene set tend to appear at the bottom of the ranked data, while a significant positive NES indicates the opposite. Cells without NES values represent non-significant enrichment (p > 0.05 after FDR correction). P: Postnatal day; SCZ: Schizophrenia.

### Enrichment Analysis for Schizophrenia-associated Genes from snRNA-seq Studies

We next assessed if the *FOXP1* gene-sets are enriched for SCZ-associated genes reported as differentially expressed in a single cell gene expression analysis of multiple cortical cell-types from SCZ patients with controls. The P7 gene-set again displayed robust enrichment for SCZ-associated genes, predominantly genes from glutamatergic excitatory neurons and, to a lesser extent, GABAergic inhibitory neurons, along with endothelial cells (Fig 4B and S8 Table). The negative enrichment observed in all the tested SCZ-associated gene-sets suggests that the downregulated genes due to *FOXP1* KO are more strongly associated with SCZ. The P47 gene-set also showed enrichment for SCZ-associated genes from almost the same cell types. However, The P7 gene-set showed stronger enrichment for SCZ-associated genes compared to P47 (Fig 4B and S8 Table). In contrast, the P0 gene-set only exhibited enrichment for SCZ-associated genes from non-neuronal cells including endothelial, oligodendrocyte progenitor cells, and microglial gene-sets (Fig 4B and S8 Table). This analysis complements our previous findings from the GWAS data, highlighting that the genes dysregulated by *FOXP1* KO at P7 and P47 overlap with genes differentially expressed within single cell-types from SCZ postmortem samples, with a stronger enrichment observed at P7.

### Functional Enrichment Analysis

We used SynGO to identify the role of *FOXP1* in synaptic processes across the postnatal stages of development. Both the P7 and P47 gene-sets were significantly enriched within many overlapping pre- and postsynaptic cellular components and biological processes, whereas the P0 gene-set was not enriched within any of these terms (S9 Table). The significant cellular components for both P7 and P47 included *postsynaptic density*, *integral components of pre- and post-synaptic membrane* and *presynaptic active zone membrane* (Fig 5 and S9 Table). The significant biological processes for both P7 and P47 included those within synapse organization (e.g. *synapse assembly*), the pre-synapse (e.g., *presynaptic vesicle exocytosis* and *regulation of presynaptic membrane potential*), the post-synapse (e.g., *regulation of postsynaptic cytosolic calcium levels* and *regulation of postsynaptic membrane neurotransmitter receptor levels*) and synaptic signaling (e.g., *chemical synaptic transmission*). We mapped genes that were both associated with SCZ and differentially expressed to enriched SynGO terms. At P7 and P47, 10 and 26 such genes mapped to these synaptic locations and functions, respectively (Fig 5).

**Fig 5.**
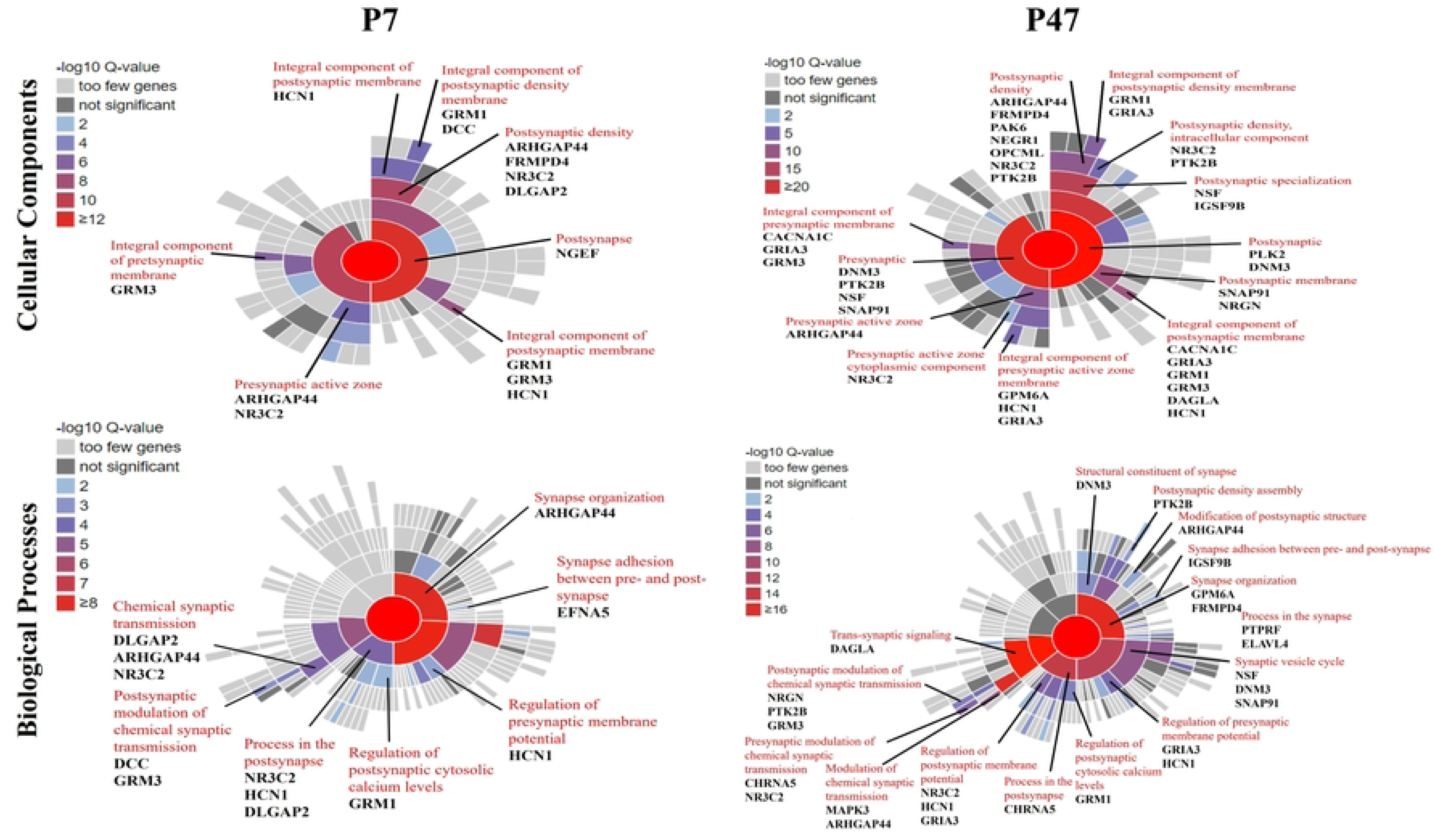
SynGO cellular component (CC) and biological processes (BP) enrichment analyses of FOXP1 DEGs identified at postnatal stages of development (P7 and P47). Sunburst plot showing enriched CC OR BP terms based on the synapse-specific SynGO database annotation. The color encodes the significance of the enriched q value. Genes involved in the significant SynGO terms and located within genome-wide significant loci for SCZ are highlighted. P: Postnatal day.

### Cell-type Enrichment Analysis

As our timepoints covered multiple stages of development, we investigated if our *FOXP1* gene-sets were enriched in individual cell types from human prenatal FC [48] and from central and peripheral nervous system of adolescent mouse brain[49]. The *FOXP1* gene-set from P0 was enriched for genes expressed in cycling progenitor cells and glutamatergic excitatory neurons within prenatal FC (S10 Table). In both the prenatal and postnatal cortex, P7 and P47 *FOXP1* gene-sets were enriched mainly within glutamatergic excitatory neurons, with the P47 gene-set also showing enrichment in GABAergic inhibitory neurons (S10 and S11 Tables).

### Trans Expression Quantitative Trait Loci Analysis

We hypothesized that genetic variation at *FOXP1* (associated with SCZ in GWAS) could influence the expression of a downstream gene, mediated through *FOXP1*’s role as a transcription factor. This would be a *trans* expression quantitative trait loci (eQTL) effect and evidence of two risk genes (i.e., *FOXP1* and a downstream gene) functioning within a putative risk pathway. To reduce the number of possible tests of target genes, we limited DEGs derived from gene expression analysis to only those genes among the 682 SCZ-risk genes reported in the latest GWAS (S12 Table) [12]. Out of the 1,697 DEGs identified across P7 and P47 developmental stages, 66 are located with genome-wide significant loci for SCZ (S13 Table). We took the rs60135207 SNP at *FOXP1,* which was associated with SCZ at genome-wide significant levels, and investigated its association with the expression levels of these 66 genes using eQTL data obtained from the Genotype-Tissue Expression (GTEx) project (https://gtexportal.org/home/) [52]. We detected a *trans* eQTL for this SCZ risk SNP at *FOXP1* with the expression of the SCZ risk gene PPIP5K1 in the cerebellum (*P*-value < 4.50E-04; S14 Table). Specifically, the T allele of rs60135207 at *FOXP1* is associated with both increased SCZ risk and increased expression of *PPIP5K1* in the cerebellum.

### Analysis of FOXP1 in Cortical Progenitor Cells

We also investigated the role of *FOXP1* in SCZ within progenitor cells during early developmental stages by analyzing prenatal data corresponding to the second trimester of human fetal development. The datasets used in this study were from progenitor cell populations, specifically embryonic mouse cortical NSCs and human cortical bRGCs derived from 3D brain organoids. During cortical development, the RGCs serve as neural progenitor cells at the ventricular zone. While migrating toward the cortical plate, they differentiate into neurons, astrocytes, and oligodendrocytes [53,54].

As a result of *FOXP1* loss of function, there were 1,075 DEGs in NSCs (E14.5) (611 upregulated and 464 downregulated) and 869 DEGs in bRGCs (362 upregulated and 507 downregulated). Comparative analysis revealed 85 DEGs common to both models, of which 39 displayed concordant expression changes (25 downregulated and 14 upregulated). Both *FOXP1* gene-sets from the two cellular models were enriched for genetic risk for SCZ (NSCs: *B* = 1.35, *p* = 1.63E-05; bRGCs: *B* = 1.37, *p* = 4.65E-06; S4 Table). These results remained significant when we omitted genes present in either the P7 or P47 gene-sets (S4 Table).

The NSC and bRGC gene-sets were both enriched in overlapping SynGO cellular compartments and biological processes, several of which were also enriched for the P7 and P47 gene-sets previously (e.g., *presynaptic active zone* and *postsynaptic density membrane, and synapse assembly* and *transsynaptic signaling*; S9 Table). Genes in significant SynGO terms and located within genome-wide significant loci for SCZ are detailed in Fig 6.

**Fig 6.**
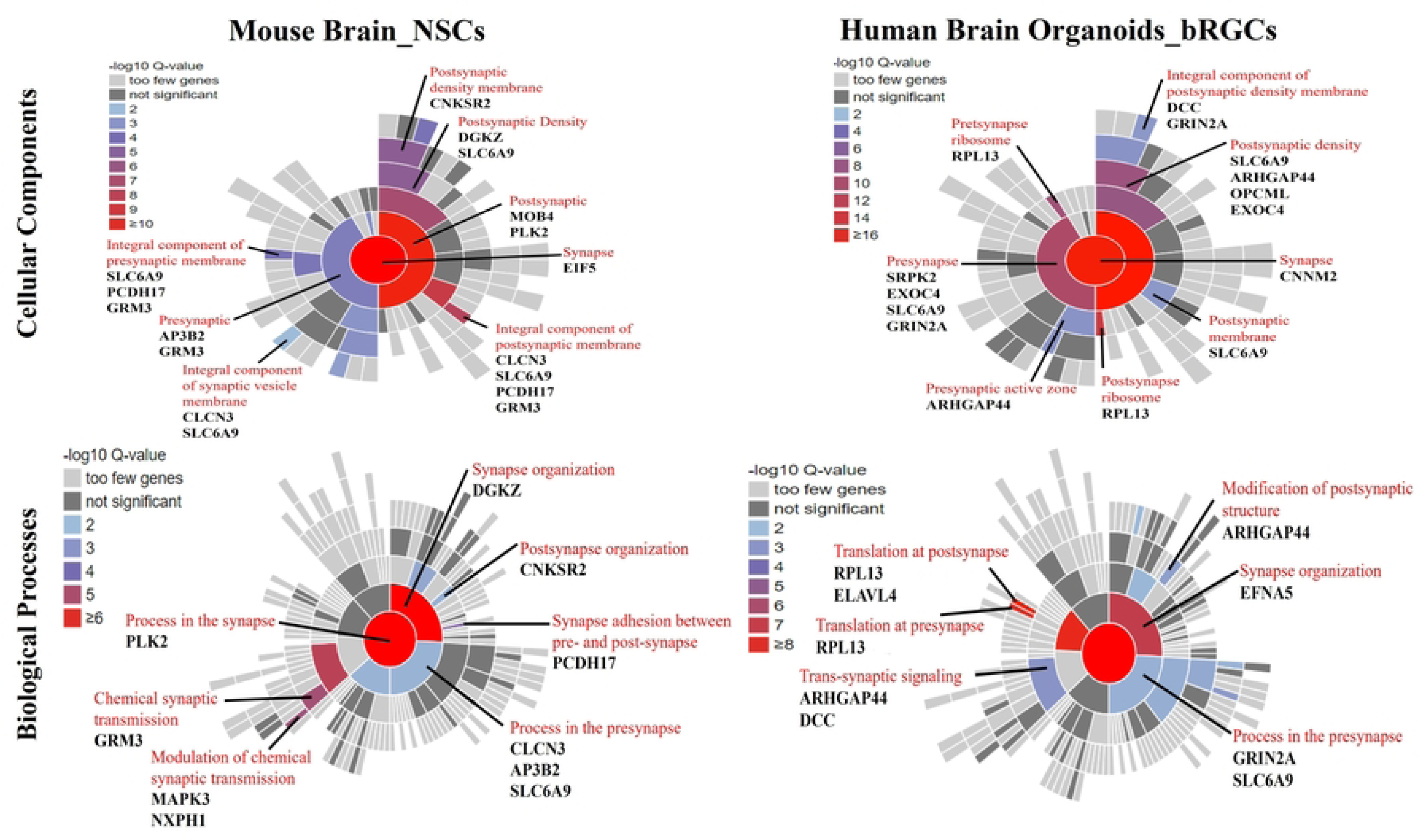
SynGO cellular component (CC) and biological processes (BP) enrichment analyses of FOXP1 DEGs identified at prenatal stages of development (second trimester). Sunburst plot showing enriched CC or BP terms based on the synapse-specific SynGO database annotation. The color encodes the significance of the enriched q value. Genes involved in the significant SynGO terms and located within genome-wide significant loci for SCZ are highlighted. NSCs: Neural stem cells; bRGCs: basal radial glial cells.

*FOXP1* gene-sets from both cellular models were tested in scRNA-seq gene expression data from human prenatal FC (S10 Table)[48]. The NSC-derived gene set was enriched within endothelial cells, oligodendrocyte precursor cells, and radial glial cells. The bRGC-derived gene set was enriched within radial glial cells (as expected), intermediate progenitors, glutamatergic excitatory neurons, and GABAergic inhibitory neurons (S10 Table).

We did not detect a *trans* eQTL for the SCZ risk SNP at *FOXP1* affecting the expression of any genes (*n*=70) altered by *FOXP1* loss of function and located within genome-wide significant loci for SCZ and in these two cellular models (S15 Table).

## Discussion

To investigate the developmental trajectory of *FOXP1*-regulated genes and their contribution to SCZ, we analyzed transcriptomic data derived from cortical cells of mouse and human models of *FOXP1* loss-of-function across different developmental stages and integrated it with human data from genetic association studies. This analysis, encompassing models spanning key developmental stages from fetal development to adolescence, revealed the dynamic nature of *FOXP1*-regulated gene sets. *FOXP1* regulates distinct gene-sets at various points in development and it is those gene-sets that are expressed at timepoints that map to the second trimester, early childhood and adolescence, but not third trimester, that are enriched for genetic risk for SCZ. These genes are expressed in different cortical cell types and involved in various synaptic functions. This highlights the value of considering developmental context when investigating the role of *FOXP1* in the pathophysiology of SCZ.

Time-course gene expression analysis for neocortical tissues across prenatal and postnatal stages identified 1,065 significant genes, where the effect of *FOXP1*-KO on their expression differed over time. Of these, 763 overlapped with the DEGs identified from the pairwise comparisons between *FOXP1*-KO and WT groups at each developmental stage. Some genes demonstrated stage-specific differential expression, while others exhibited altered expression across multiple stages. Clustering analysis of these genes revealed distinct expression patterns, grouping them into 20 clusters based on their patterns of expression across the postnatal stages. Among the genes in the clusters that exhibited distinct expression patterns between the *FOXP1*-KO and WT groups across one or more developmental stages, three (*PLK2*, *CACNA1I*, and *NEK1*) are located with genome-wide significant loci for SCZ. The protein encoded by *PLK2* is a member of the polo-like kinase (PLK) family of serine/threonine protein kinases, crucial for normal cell division [55]. During brain development, PLK2 is expressed in the cortical plate of embryos and its expression is upregulated by BDNF signaling, promoting dendritic growth in immature cortical neurons [56]. It has been shown to respond to synaptic activity, playing a crucial role in spine formation and the regulation of synaptic homeostasis [57,58]. Similarly, NEK1, another serine/threonine kinase, is involved in cell cycle control, ciliogenesis, and the DNA damage response [59–61]. The *CACNA1I* gene encodes CaV3.3, a T-type voltage-gated calcium channel that regulates neuronal excitability and rhythmic activity within neuronal circuits [62]. Functional studies of a *CACNA1I* variant suggest that reduced CaV3.3 activity may confer protection against SCZ by decreasing excitability within the thalamic reticular nucleus [63]. These findings underscore the significance of *FOXP1* in regulating the expression of genes critical for neuronal development and synaptic function across different stages of brain development, providing valuable insights into the potential molecular mechanisms underlying the pathophysiology of SCZ.

Pairwise comparisons across P0, P7, and P47 revealed that *FOXP1* regulates gene expression in a stage-dependent manner during development. While some DEGs were common across two stages of development, and a few were consistently altered across all three stages, the majority of the DEGs exhibited stage-specific changes, indicating that *FOXP1*’s transcriptional impact dynamically changes throughout development. These findings emphasize the importance of developmental timing in understanding *FOXP1*’s potential contribution to SCZ. Using sLDSC, we observed the strongest enrichment for SCZ risk within *FOXP1*-regulated genes was at the P7 stage, followed by P47, with no significant enrichment detected at the earlier P0 stage. In addition to examining SCZ risk indexed by GWAS, we investigated whether *FOXP1*-related gene sets are enriched for SCZ-associated genes identified by gene expression analysis of cortical cell types from SCZ patients. Like the GWAS-based analysis before, the strongest enrichment was observed at the P7 stage and specifically within glutamatergic excitatory neurons. This finding, consistent with the predominant expression of *FOXP1* in these neuronal subtypes [24], suggests a link between *FOXP1*-mediated gene regulation and the development of SCZ in this specific neuronal population at this stage of development. SynGO enrichment analysis was specifically performed within gene-sets exhibiting significant enrichment for genes associated with SCZ at P7 and P47. This analysis highlighted *FOXP1*’s role in postnatal synaptic processes, with P7 and P47 gene-sets significantly enriched within multiple SynGO terms, important for synaptic connectivity and function.

Among these genes, *HCN1,* encodes a voltage-gated potassium/sodium channel that is a main contributor to hyperpolarization-activated cation current. In addition to its role in regulating neuronal excitability, *HCN1* has also been found in animal studies to play a significant role in rhythmic activity, synaptic plasticity [64]. Recent studies show that HCN1 is associated with working memory impairments in SCZ patients [65]. In addition to HCN1, other genes implicated in synaptic plasticity and neuronal excitability, such as *GRM1* [66] and *GRM3* [67], also emerge as potential contributors to cognitive dysfunction in SCZ. *GRM1* and *GRM3* encode metabotropic glutamate receptors (mGluRs), members of the G-protein coupled receptor (GPCR) superfamily, which play a key role in neurotransmitter signaling within the brain [68]. *FRMPD4* is a positive regulator of dendritic spine morphogenesis and density through interaction with PSD-95 [69]. It is essential for maintaining excitatory synaptic transmission through interaction with mGluR1/5 [70]. Recent studies have shown that polymorphisms in the human *FRMPD4* gene are associated with sex differences in SCZ, and mutations in *FRMPD4* can cause X-linked ID [71,72]. At P7, two genes associated with SCZ, *DLGAP2* and *NGEF*, are linked to dendritic spine density and synaptic function. DLGAP2 is also a scaffolding protein, directly interacts with PSD-95. *De novo* mutations have been reported in *DLGAP2* gene in SCZ patient cohorts [73], further emphasizing its importance in the pathology of the disorder.

The most recent GWAS of SCZ identified 682 genes within 287 genome-wide significant loci [12]. We identified a trans eQTL effect of a SNP in *FOXP1* on the expression of one of these risk loci. Disruption of *FOXP1* at P47 stage resulted in reduced expression of *PPIP5K1*. *PPIP5K1* encodes a dual functional inositol kinase [74]. This enzyme regulates inositol phosphate metabolism, a pathway increasingly implicated in SCZ pathophysiology [75,76], suggesting *FOXP1*’s role in modulating SCZ-associated molecular mechanisms.

We also explored *FOXP1* function in SCZ pathogenesis during second trimester-equivalent prenatal development by analyzing gene expression in mouse NSCs (E14.5) and human bRGCs. Both models showed significant dysregulation of gene expression in response to *FOXP1* loss, with enrichment for genes associated with SCZ. SynGO analysis showed that these genes are involved in a wide range of synaptic functions. The mapping of SCZ-associated genes to SynGO-enriched terms demonstrated both overlapping and unique gene sets between prenatal and postnatal stages, indicating stage-specific risk factors for SCZ. Among these, one gene, *SLC6A9*, was common between the two cellular models analyzed in the prenatal stage. *SLC6A9* encodes the GLYT1 glycine transporter, which is responsible for maintaining low levels of glycine, an N-methyl-D-aspartate receptor (NMDAR) co-agonist, in the synaptic cleft. This suggests that *SLC6A9* may play a role in the development of NMDAR hypofunction, which has been implicated in SCZ [77]. *ARHGAP44*, a common gene between the prenatal and postnatal stages, is a synaptic Rho-GAP that binds to the SHANK3 protein, which is involved in dendritic spine formation and synaptic plasticity [78]. The SHANK family is closely associated with ASD and SCZ, and its interaction with SHANK3 points to a possible involvement of *ARHGAP44* in neuropsychiatric conditions [79].

A limitation of the study is its reliance primarily on RNA-seq data from mouse models. While this approach provides valuable insights, it may not fully capture the complexity of *FOXP1* function across species. Secondly, the study focused only on specific developmental stages where data were available and that did not include a timepoint equivalent to adulthood in humans. Thirdly, the mouse transcriptomic data corresponding to the third trimester, early childhood, and adolescence in humans are generated from bulk-tissue RNA-seq and lack cell-type resolution. In contrast, the data representing the second trimester were derived from progenitor cells, generated either through snRNA-seq or by preforming *FOXP1* knockdown specifically in NSCs *in vitro*. This reflects distinct and specific cell types from those analyzed in the other studied stages.

This study leverages *FOXP1*’s association with SCZ and gene expression data from biological models of *FOXP1* to demonstrate that *FOXP1* plays a dynamic role in regulating gene expression across development, with a significant impact on genes associated with SCZ risk, particularly during the second trimester, early childhood and adolescence. Our findings highlight the importance of considering the dynamic nature of brain development when investigating the genetic underpinnings of SCZ. By identifying key genes and pathways impacted by *FOXP1* loss, including those involved in synaptic function, this study provides insights into the molecular mechanisms underlying *FOXP1*’s contribution to SCZ susceptibility.

## Funding

This work was funded by grants from the University of Galway, Ireland (Hardiman Research Scholarship #128936 to DA).

## Conflict of Interest

The authors declare no conflict of interest.

## Acknowledgements

The authors would like to thank Dr. Aodán Laighneach (University of Galway) for his valuable input on the methodology used in this study.

## Suplemmentary Information

S1 Table: Lists of DEGs (Ruzicka *et al.,* 2024) used for competitive GSEA.

S2 Table: Background Gene Lists Used for SynGO Analysis.

S3 Table: Significant genes identified in time-course gene expression analysis

S4 Table: sLDSC Analysis Results of FOXP1 Gene-Sets Using GWAS Data for SCZ and Control Phenotypes

S5 Table: Gene expression clusters of genes identified in time-course gene expression analysis

S6 Table: Genes Identified from Time-Course Gene Expression Analysis and Located within Genome-Wide Significant Loci of SCZ

S7 Table: Significant DEGs identified in pairwise gene expression analysis

S8 Table: GSEA Analysis of FOXP1 gene-sets Using Data on snRNA-seq

S9 Table: SynGO Analysis for FOXP1 Gene-sets

S10 Table: EWCE analysis of FOXP1 Gene-Sets in the Cameron et al., 2023 scRNA-seq Data for FC Brain Region from Prenatal Brain.

S11 Table: EWCE analysis of FOXP1 Gene-Sets in the Zeisel et al. 2018 scRNA-seq Data for FC Brain Region from postnatal Brain.

S12 Table: SCZ Risk Genes reported in the Latest GWAS for SCZ (Trubetskoy et al., 2022)

S13 Table: List of Genes Overlapping between DEGs (S7 Table) and Genome-wide Significant Loci for SCZ Reported in the Latest SCZ GWAS (S12 Table)

S14 Table: eQTL Analysis for the LD-Independent FOXP1 SNP on Genome-wide significant loci for SCZ within P7 and P47 DEGs and Cell-Type Specific Expression in Different Brain Tissues Based on the GTEx Dataset.

S15 Table: eQTL Analysis for the LD-Independent FOXP1 SNP on Genome-wide significant loci for SCZ within NSCs_E14.5 and Human Brain organoid_bRGCs DEGs and Cell-Type Specific Expression in Different Brain Tissues Based on the GTEx Dataset.

## Notes

### Competing Interest Statement

The authors have declared no competing interest.

